# Assessing the utility of marine filter feeders for environmental DNA (eDNA) biodiversity monitoring

**DOI:** 10.1101/2021.12.21.473722

**Authors:** Gert-Jan Jeunen, Jasmine S. Cane, Sara Ferreira, Francesca Strano, Ulla von Ammon, Hugh Cross, Robert Day, Sean Hesseltine, Kaleb Ellis, Lara Urban, Niall Pearson, Pamela Olmedo-Rojas, Anya Kardailsky, Neil J. Gemmell, Miles Lamare

## Abstract

Aquatic environmental DNA (eDNA) surveys are transforming how we monitor marine ecosystems. The time-consuming pre-processing step of active filtration, however, remains a bottleneck. Hence, new approaches omitting active filtration are in great demand. One exciting prospect is to use the filtering power of invertebrates to collect eDNA. While proof-of-concept has been achieved, comparative studies between aquatic and filter feeder eDNA signals are lacking. Here, we investigated the differences among four eDNA sources (water; bivalves; sponges; and ethanol in which filter-feeding organisms were stored) along a vertical transect in Doubtful Sound, New Zealand using three metabarcoding primers (fish (16S); MiFish-E/U). While concurrent SCUBA diver observations validated eDNA results, laboratory trials corroborated in-field bivalve eDNA detection results. Combined, eDNA sources detected 59 vertebrates, while divers observed eight fish species. There were no significant differences in alpha and beta diversity between water and sponge eDNA and both sources were highly correlated. Vertebrate eDNA was detected in ethanol, although only a reduced number of species were detected. Bivalves failed to reliably detect eDNA in both field and mesocosm experiments. While additional research into filter feeder eDNA accumulation efficiency is essential, our results provide strong evidence for the potential of incorporating sponges into eDNA surveys.

## 1 INTRODUCTION

The incorporation of species detection through bulk DNA obtained from environmental samples (environmental DNA, eDNA) is revolutionizing the way researchers monitor the marine biome (Bowers et al., 2021; Ficetola et al., 2008). By inferring species presence and absence indirectly through molecular approaches, eDNA surveys do not rely on visual observations, thereby simplifying species detection within inaccessible environments (Fediajevaite et al., 2021). In addition, the complexity of the DNA signal contained within environmental samples facilitates the detection of a broad range of taxonomic groups, and a larger number of individual species, than can be achieved using traditional monitoring (Jeunen et al., 2020; Kelly et al., 2017; Kim et al., 2019; Thomsen et al., 2012). The potential of eDNA surveys has particular appeal and application for marine conservation efforts, where comparative experiments show eDNA routinely outperforms traditional monitoring (Afzali et al., 2020; Bakker et al., 2017; Kelly et al., 2017; Port et al., 2016; Stat et al., 2019; Thomsen et al., 2012, 2016; Valdivia-Carrillo et al., 2019).

The lack of visual species observation, however, has warranted a thorough investigation into the accuracy of aquatic eDNA surveys in the marine environment (Kelly et al., 2014; Parsons et al., 2020; von Ammon et al., 2019). Thus far, eDNA signals have been positively correlated with species presence (Collins et al., 2018; Jo et al., 2017; Murakami et al., 2019; Saito & Doi, 2021; Wood et al., 2020), and high spatial and temporal resolutions have been observed (Berry et al., 2019; Djurhuus et al., 2020; Jeunen et al., 2019b, 2019a; O’Donnell et al., 2017; Port et al., 2016; Sigsgaard et al., 2017; Stoeckle et al., 2017; Uthicke et al., 2018). However, increased spatial and temporal sampling effort is required to detect the marine community within an area and minimize the risk of false-negative species detections (Griffin et al., 2020; Pinfield et al., 2019; Wilcox et al., 2016). Furthermore, increased sampling effort is hindered by the long process of vacuum filtration to capture eDNA from water and the continuous eDNA degradation starting from sample collection until processing (Curtis et al., 2021; Hinlo et al., 2017).

Recently, alternative options to active filtration have been explored to circumvent the need for the time-intensive process of vacuum filtration and facilitate the implementation of large-scale eDNA surveys (Bessey et al., 2021; Bowers et al., 2021; Kirtane et al., 2020; Mariani et al., 2019; Turon et al., 2020; Verdier et al., 2021; Yamahara et al., 2019). Passive filtration strategies, whereby filter membranes (Bessey et al., 2021) and other DNA-absorbing matrices (Kirtane et al., 2020) are submerged in the environment, have successfully captured eDNA in seawater. Furthermore, passive filtration strategies have shown to perform equally to active filtration (Bessey et al., 2021). However, the necessary submergence time (>8 hours) and physical limits of filter membranes to passively absorb eDNA might hinder its implementation (Bessey et al., 2021).

To avoid long submergence times and potentially complicated deployment and retrieval of passive filtration devices, organisms with a filter-feeding strategy might be utilized if they naturally accumulate eDNA (Mariani et al., 2019). These organisms may be especially useful for eDNA surveys, because they can filter large volumes of water - sponges filter up to 900 times their volume of sea water per day (Gökalp et al., 2020), while bivalves can filter over 200 L per day (Berge et al., 2006). While filter feeders are scattered across the tree of life (Hickman et al., 1995), only the eDNA accumulation efficiency of sponges (Mariani et al., 2019; Turon et al., 2020) and shrimp (Siegenthaler et al., 2019) has, thus far, been investigated. Information on other filter-feeding taxa, development of optimal eDNA extraction protocols, and comparative studies determining the comparability between aquatic and filter feeder eDNA are currently lacking. As both protocol (Deiner et al., 2018; Jeunen et al., 2018) and substrate choice (Koziol et al., 2019; Turner et al., 2015) are known to affect downstream eDNA analysis, additional research into filter feeder eDNA efficacy is needed to determine their suitability for eDNA surveys.

In this study, we compared eDNA signals obtained from (i) water samples, (ii) bivalve gill-tissue dissections, (iii) sponge tissue, and (iv) the ethanol in which filter-feeding specimens were stored. This comparison will allow us to assess the viability of including marine filter-feeding organisms into eDNA surveys and, thereby, facilitating large-scale applications. Furthermore, we validated eDNA detections by concurrent SCUBA diver observations. Additionally, as this experiment is, to the best of our knowledge, the first to investigate the use of bivalve gill-tissue for eDNA detection, a mesocosm experiment was set up to corroborate in-field mussel eDNA detection results. We address four questions:

1. Are eDNA signals obtained from filter-feeding organisms and water samples comparable?
2. Does the eDNA signal obtained from filter-feeding organisms display identical biodiversity patterns along a stratified vertical transect compared to aquatic eDNA?
3. Can eDNA signals accumulated within filter-feeding organisms be obtained from the ethanol they are stored in?
4. How do eDNA surveys compare against traditional diver surveys in detecting vertebrate diversity?

## 2 RESULTS

### 2.1 Diver survey

Our 1-hour SCUBA diver survey by three experienced marine scientists in Doubtful Sound (Figure 1a) identified a total of eight fish species, all of which could be identified to species-level based on visual morphological characteristics (Supplement 1). The survey consisted of divers swimming 50 m horizontal transects at 15, 10, and 5 m depth strata. Conditions were ideal for diver surveys, with good horizontal visibility (10 to 15 m) and an open rock wall. Diver observations were grouped for all sampling depths, excluding the persistent low-salinity layer (LSL, Figure 1b) where no observations were recorded due to high turbidity and low visibility (Gibbs et al., 2000). Abundance of fish was classified by divers in three categories: ‘abundant’, ‘common’, and ‘rare’. Butterfly perch (*Caesioperca lepidoptera*) and spotty wrasse (*Notolabrus celidotus*) were encountered as ‘abundant’. ‘Common’ species included the banded wrasse (*Notolabrus fucicola*), Jock Stewart (*Helicolenus percoides*), oblique triplefin (*Forsterygion maryannae*), and common triplefin (*Forsterygion lapillum*). Blue cod (*Parapercis colias*) and scarlet wrasse (*Pseudolabrus miles*) were only sighted once during the diver survey and, therefore, noted down as ‘rare’ sightings. No sharks, mammals, and birds were seen by the divers or by researchers on the boat collecting aquatic eDNA samples during the survey period.

**Figure 1:**
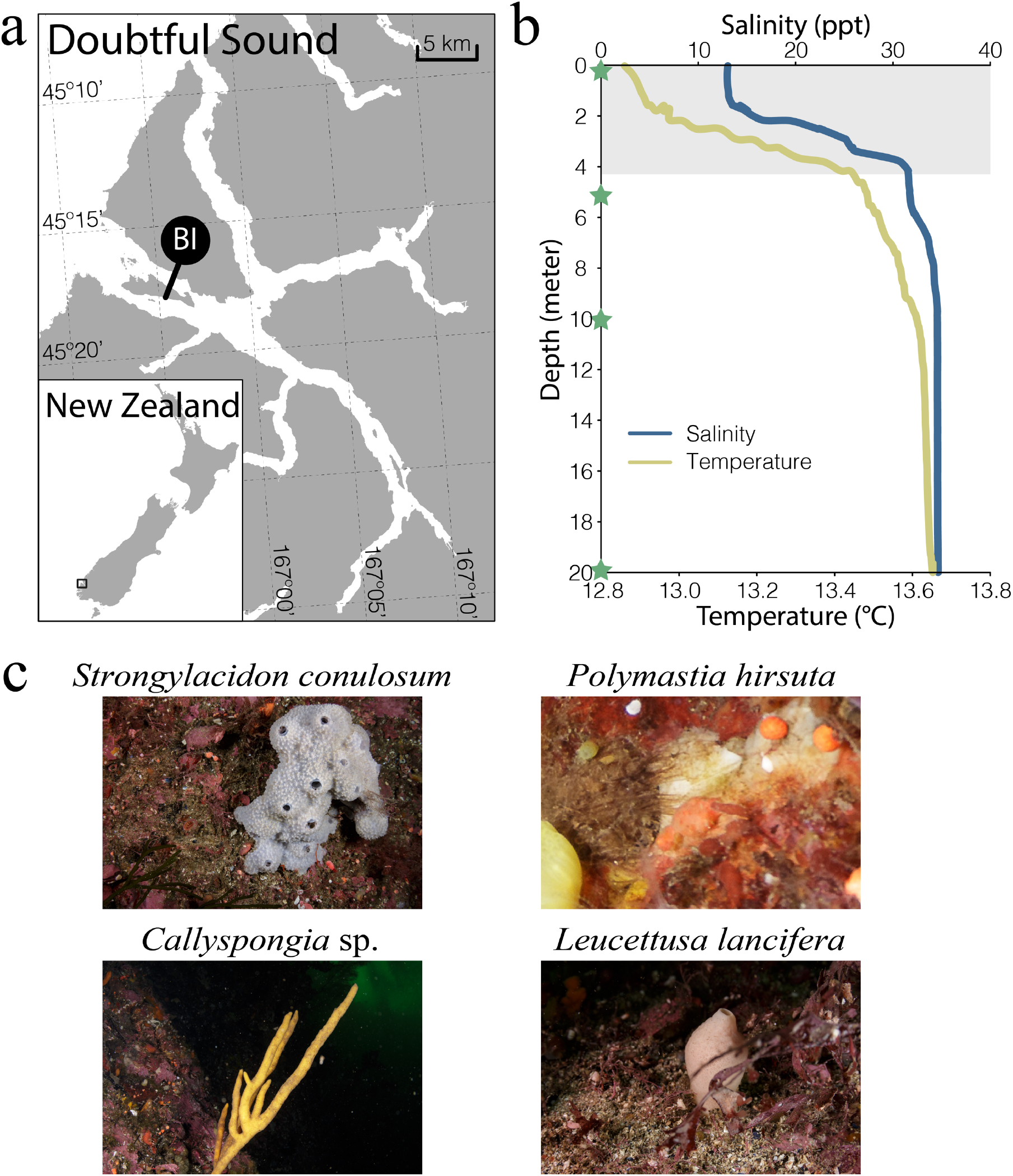
(a) Map of Doubtful Sound, New Zealand with the sample collection site Bauza Island (BI) indicated by a black circle. (b) Depth profile as measured by CTD profiler. Y-axis displays depth, while the top and bottom x-axes display salinity (blue line) and temperature (yellow line), respectively. The low-salinity layer (LSL) is indicated by the shaded gray area. Sampling depths (0, 5, 10, and 20 m) are indicated by a green star. (c) Photographs of sponges sampled during field work (photo credit: Adam Brook). Sponge identification obtained through spicule analysis was performed by Francesca Strano.

### 2.2 Morphological identification of sponge specimens

Spicule analysis revealed that four sponge taxa belonging to the classes Calcarea and Demospongiae were sampled during the diver survey (Figure 1c). *Strongylacidon conulosum* (Bergquist & Fromont, 1988; Supplement 2h-l), *Callyspongia* sp. (Duchassaing & Michelotti, 1864; Supplement 2d,e), and *Polymastia hirsuta* (Bergquist, 1968; Supplement 2f,g) were sampled at 5 m depth, while *Leucettusa lancifera* (Dendy, 1924; Supplement 2a-c) was sampled at 10 m and 20 m depth.

### 2.3 Environmental DNA survey

#### 2.3.1 Sequencing results

Filtering and quality control returned a total of 7,196,742 sequencing reads (fish (16S): 1,741,673 reads; MiFish-E: 2,808,711 reads; MiFish-U: 2,646,358 reads; Supplement 3). Samples without a single read were removed from subsequent analyses, including one mussel gill-tissue dissection (BIM_5), all ethanol samples in which mussel gill-tissue dissections were stored (E_BIM_1, E_BIM_2, E_BIM_3, E_BIM_4, E_BIM_5), and one ethanol sample in which a sponge was stored (E_BISP_10_1). Overall, eDNA samples achieved sufficient sequencing coverage, based on rarefaction curves reaching the asymptote (Supplement 4) and mean number of reads per sample ± SD: fish (16S): 36,284 ± 25,190; MiFish-E: 58,514 ± 34,979; MiFish-U: 94,432 ± 55,866. Negative control samples contained one OTU (Operational Taxonomic Unit; taxonomic ID: human) for the MiFish-U/E metabarcoding primers, which was excluded from the full dataset during quality filtering prior to statistical analysis. No reads were returned for negative control samples for the fish (16S) metabarcoding primer.

#### 2.3.2 Taxonomic diversity

After stringent quality control and OTU clustering, 126, 113, and 129 OTUs were obtained for the fish (16S), MiFish-E, and MiFish-U metabarcoding primers, respectively. Taxonomic assignment identified a total of 59 unique taxonomic IDs (fish (16S): 25; MiFish-E: 45; MiFish-U: 33). While the taxa identified by the three metabarcoding primers were observed to be similar at the order (KDI_order_ = 0.24 ± 0.08) and species level (KDI_species_ = 0.40 ± 0.12), only partial overlap of diversity was achieved between metabarcoding primers at both taxonomic levels (Supplement 5). The total number of taxonomic IDs covered 46 families within 29 orders and four classes in the phylum Chordata. The class Actinopterygii was the most abundant, consisting of 40 taxonomic IDs (67.8%) and 6,799,436 reads (94.5%), followed by Mammalia (taxonomic IDs: 11; 18.6%, reads: 286,175; 4.0%), Aves (taxonomic IDs: 5; 8.5%, reads: 93,686; 1.3%), and Chondrichthyes (taxonomic IDs: 3; 5.1%, reads: 17,445; 0.2%).

### 2.4 Aquatic, filter-feeder, and ethanol eDNA comparison

#### 2.4.1 Alpha diversity measures

Across all sampling depths, aquatic eDNA detected the largest number of taxa (54), followed by filter-feeder eDNA (31 taxa) and ethanol eDNA (20 taxa; Figure 2). The increased number of taxa detected by aquatic eDNA is attributed to eDNA signals from freshwater (e.g., *Anguilla dieffenbachii*, New Zealand longfin eel; *Oncorhynchus tshawytscha*, chinook salmon; *Galaxias brevipinnis*, kōaro or climbing galaxias; and *Gobiomorphus cotidianus*, common bully) and terrestrial organisms (e.g., *Trichosurus vulpecula*, common brushtail possum; *Ovis aries*, sheep; *Gallirallus australis*, weka; and *Petroica macrocephala*, tomtit), which are missed by filter-feeder and ethanol eDNA. However, both filter-feeder and ethanol eDNA detected taxa not recovered by aquatic eDNA (e.g., *Megaptera novaeangliae*, humpback whale; *Cottus* sp., sculpin fish; Figure 2; Supplement 3).

**Figure 2:**
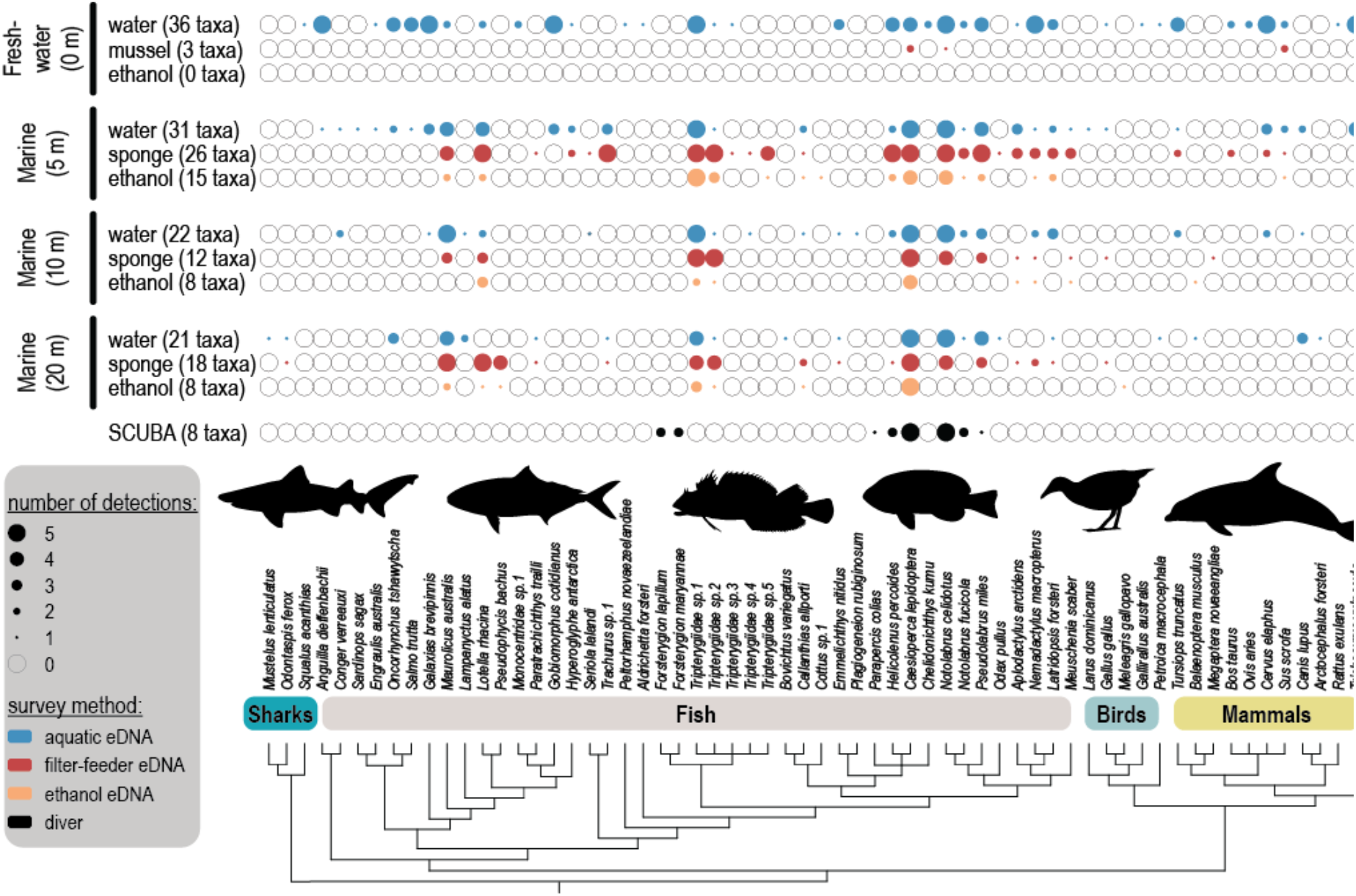
Observed diversity at Bauza Island, Doubtful Sound, New Zealand for each sampling depth (0, 5, 10, 20 m) and survey method (aquatic eDNA; mussel eDNA; sponge eDNA; ethanol eDNA; SCUBA diver survey). Filled and unfilled circles indicate taxon presence or absence. Circle diameter indicates the number of positive detections within each sampling depth and survey method (maximum 5 replicates for each treatment). Aquatic eDNA is colored in blue, filter-feeder eDNA in red, ethanol eDNA in orange, and diver detections in black.

Within each sampling depth, an increased number of taxa were detected by aquatic eDNA compared to filter-feeder and ethanol eDNA according to species accumulation curves (Figure 3a). Furthermore, significant differences in alpha diversity were observed across eDNA sources and sampling depths (Figure 3b) according to ANOVA (F_11,48_ = 38.75; *P* < 0.0001). Post hoc Tukey-Kramer revealed a significant difference in alpha diversity among aquatic eDNA and mussel and ethanol eDNA (sampling depth: 0 m), with aquatic eDNA collected in the LSL displaying the highest diversity across all sampling depths and eDNA sources, and mussel gill-tissue dissections failing to reliably detect vertebrate eDNA signals. Mussel gill-tissue dissections managed on average a single detection per sample, with three unique taxa detected across all samples, i.e., butterfly perch (*Caesioperca lepidoptera*; most abundant sequence in the overall dataset), spotty wrasse (*Notolabrus celidotus*; third most abundant sequence in the overall dataset), and pig (*Sus scrofa*; seventeenth most abundant sequence in the overall dataset). Additionally, ethanol samples in which mussel gill tissue dissections were stored failed to detect a single taxon (Figure 2; Figure 3b).

**Figure 3:**
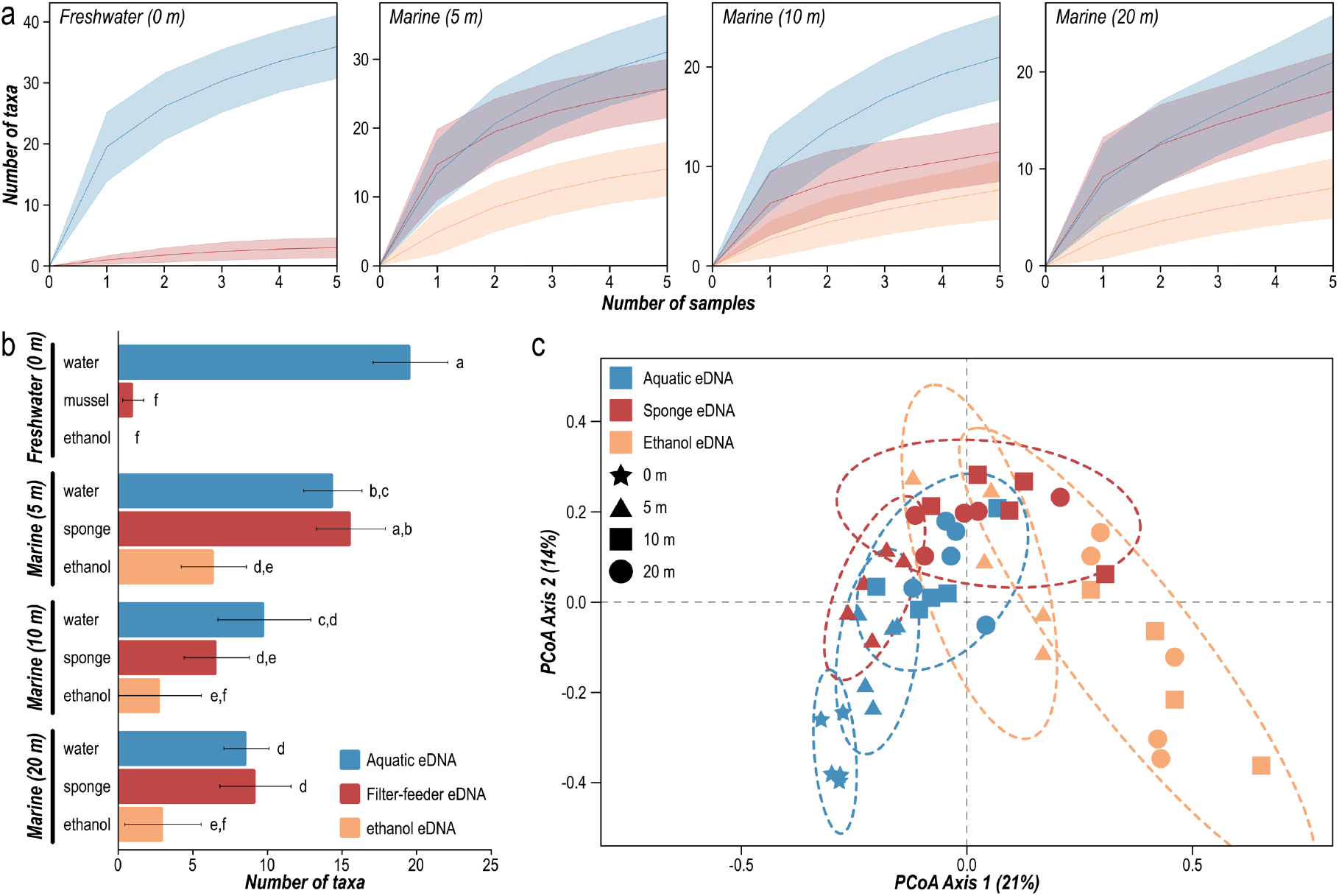
(a) Species accumulation curves per sampling depth (0, 5, 10, and 20 m), with number of samples on x-axis and number of species on y-axis. Line indicates the average value, while shaded area depicts the standard error. Aquatic eDNA is represented in blue, filter-feeder eDNA in red, and ethanol eDNA in orange. (b) The average taxonomic richness obtained per replicate for each eDNA source (aquatic, filter-feeder, and ethanol eDNA) at each sampling depth (0, 5, 10, and 20 m). Error bars show 95% confidence intervals. Significant differences as indicated by post hoc Tukey-Kramer are depicted by a lower-case letter. (c) Principal Coordinates Analysis (PCoA) depicting similarity in community composition based on eDNA taxonomic incidence (Jaccard index; presence-absence), with the primary x-axis explaining 21% of variation seen in the dataset and secondary y-axis explaining 14% of variation. Samples collected in the low-salinity layer (LSL, 0 m) are indicated by a star, 5 m with a triangle, 10 m with a square, and 20 m with a circle. Ellipses surrounding each group of samples represent 95% confidence intervals.

No significant difference was observed between aquatic and sponge eDNA for samples collected at 5 m, 10 m, and 20 m deep (Figure 3b). Pairwise significant differences in alpha diversity (*P* < 0.05), however, were observed between these depths irrespective of eDNA source (aquatic or sponge). Furthermore, while vertebrate eDNA signals were successfully retrieved from ethanol samples in which sponge specimens were kept, alpha diversity measures were significantly reduced compared to aquatic and sponge eDNA (Figure 3b).

#### 2.4.2 Beta diversity measures

Significant differences in community composition were observed across depth and eDNA source according to PERMANOVA (*F*_9,39_ = 5.2098; *P* < 0.001). PERMDISP analysis revealed no significant differences in dispersion (*F*_9,39_ = 1.2692; *P* = 0.3). Further statistical evidence for the partitioning of samples between depths and eDNA sources was confirmed by ordination analysis (PCoA analysis; Figure 3c), which revealed that the observed community composition differences were mostly driven by the number of species detected. Separation of samples on the primary axis explaining 21% of the variation was observed between ethanol eDNA samples and sponge and aquatic eDNA samples (Figure 3c). Secondarily, community composition differences were driven by the vertical zonation structure present in Doubtful Sound, with samples separating based on depth along the secondary axis explaining 14% of the variation (Figure 3c).

#### 2.4.3 Read abundance correlation between eDNA sources

Linear regression analysis showed a significant correlation in square-root transformed read abundance for detected diversity between all three eDNA signal sources (water – sponge: R^2^ = 0.7763, *P* < 0.001; sponge – ethanol: R^2^ = 0.8886, *P* < 0.001; water – ethanol: R^2^ = 0.7029, *P* < 0.01; Figure 4a-c). When excluding terrestrial and freshwater eDNA signals, correlation in read abundance significantly increased between all three eDNA signal sources (water – sponge: R^2^ = 0.8478, *P* < 0.001; sponge – ethanol: R^2^ = 0.8929, *P* < 0.001; water – ethanol: R^2^ = 0.7766, *P* < 0.01; Figure 4d-f).

**Figure 4:**
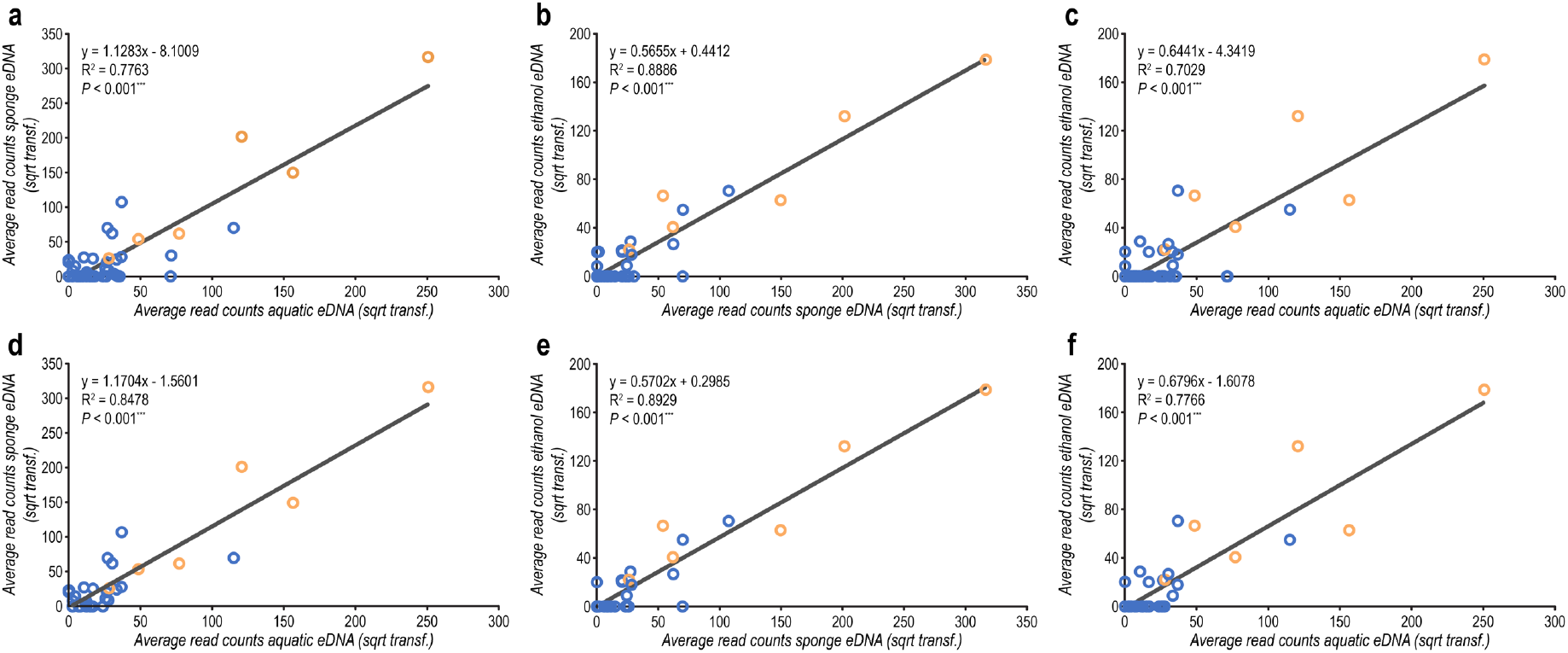
Correlations between eDNA signal sources. Scatterplots showing lines of best fit and linear regressions on the total dataset (a-c) and exclusion of terrestrial and freshwater eDNA signals (d-f) for square-root transformed average read count. Panel (a) and (d) display the correlation between aquatic and sponge eDNA. Panel (b) and (e) display the correlation between sponge and ethanol eDNA. Panel (c) and (f) display the correlation between aquatic and ethanol eDNA. Taxa detected by diver survey are depicted as orange circles.

### 2.5 eDNA and diver survey diversity comparison

Excluding samples collected in the LSL (no diver observations were made due to poor visibility), a total of 54 taxa were detected in the marine water layer (Figure 2; Supplement 3), with aquatic eDNA detecting the largest number of taxa (44; 81%), followed by sponge eDNA (31; 57%), ethanol eDNA (20; 37%), and diver survey (8; 15%). Although eDNA detected a significantly larger number of taxa compared to diver observations, only partial overlap in species detection was observed between survey methods and eDNA sources (Figure 5a). The diver survey detected three fish species not picked up by eDNA, including blue cod (*Parapercis colias*; sighted once and noted as ‘rare’ by diver), oblique triplefin (*Forsterygion maryanne*; noted as ‘common’ by diver), and common triplefin (*Forsterygion lapillum*; noted as ‘common’ by diver). The five taxa detected by diver and eDNA constituted 63.3% ± 8.3% of reads (Figure 5b). While the two triplefin species were not detected by eDNA, sequencing data included reads assigned to the triplefin family (Tripterygiidae), with species assignment not feasible due to a lack of reference sequences. Including reads assigned to the family Tripterygiidae, proportion of reads assigned to taxa observed by diver constituted 80.7% ± 6.3% of the total number of reads.

**Figure 5:**
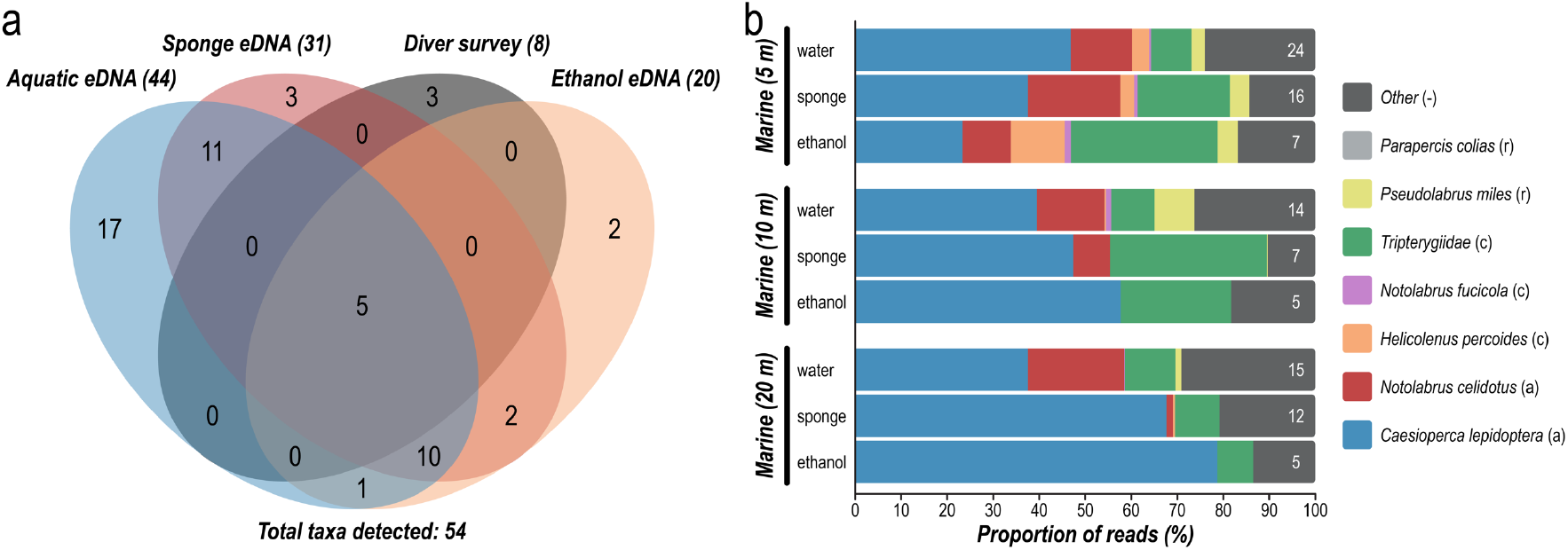
(a) Venn diagram showing overlap in species detection among the four survey methods and eDNA sources. Numbers represent the number of species detected summed over all replicates. Total number of detected species is indicated for each method between brackets alongside the name. (b) Proportion of eDNA reads assigned to each of the eight detected species by diver (colored) and remaining number of species only detected by eDNA (grey). Number within the grey bar represents the number of additional taxa detected by eDNA summed over five replicates. Abundance of diver-detected species is indicated between brackets with: ‘a’ – abundant; ‘c’ – common; and ‘r’ – rare.

### 2.6 Mesocosm experiment

Mussel and ethanol eDNA only managed to detect one of the seven fish species in the mesocosm across all replicates (Figure 6), i.e., *Nemadactylus macropterus* (terakihi). One additional fish species was detected in two replicates, i.e., *Notolabrus celidotus* (spotty wrasse). However, active filtration eDNA experiments undertaken concurrently with mussel sampling detected all but one of the fish species (*Acanthoclinus fuscus*) present in the mesocosm (Jeunen and von Ammon et al., *in prep*.). All active filtration eDNA detections were consistent across the five replicates, with exception of *Latris lineata*, which was only detected in a single replicate. Additionally, negative in-field eDNA detection results were also obtained for another bivalve species, i.e., New Zealand cockle (*Austrovenus stutchburyi*) in a separate in-field experiment (Supplement 6), in which Sanger sequencing of 16S-fish amplicons revealed sequences to be most likely derived from uncultured bacterial DNA.

**Figure 6:**
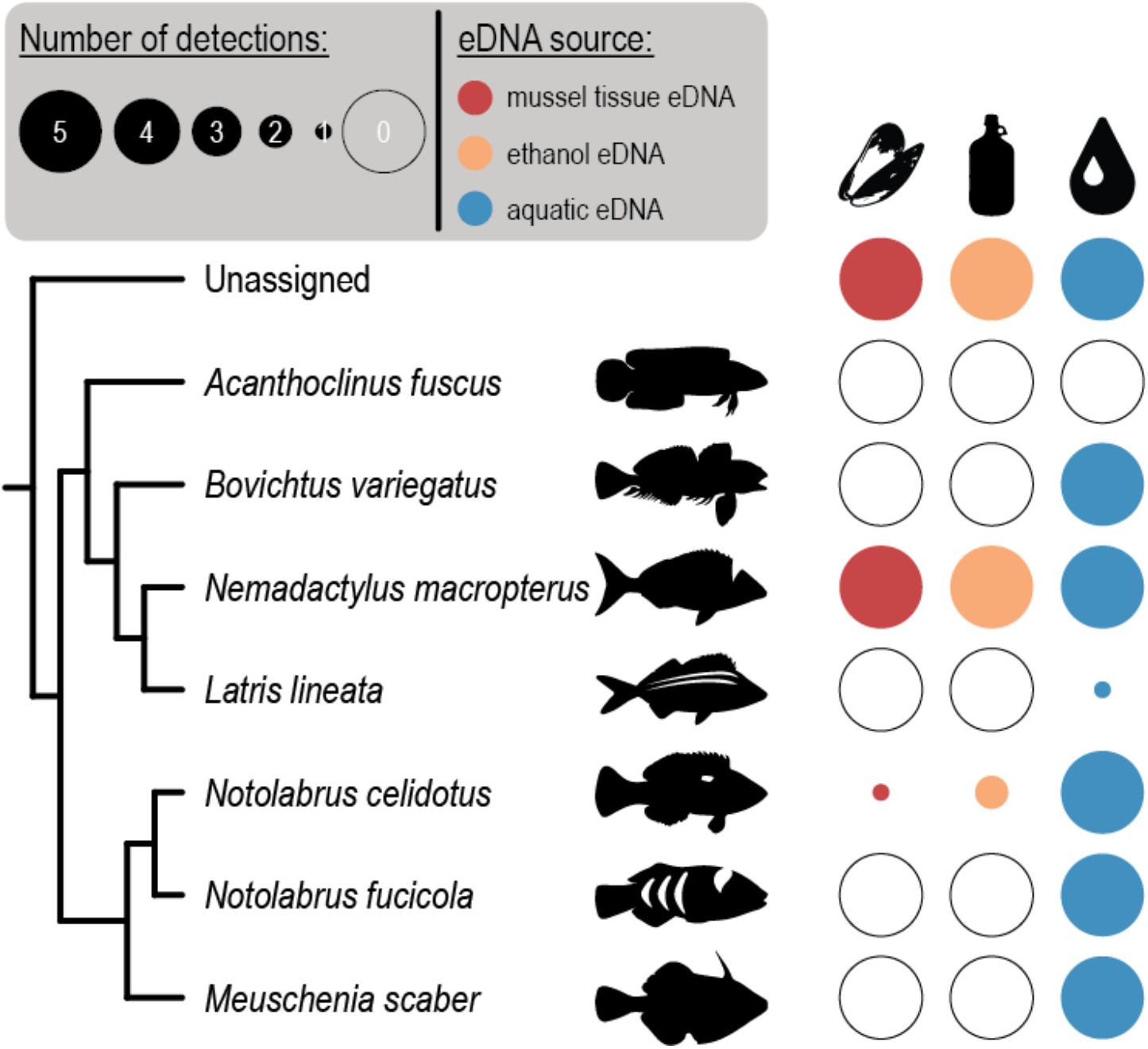
Species detection of mussel (red) and ethanol eDNA (orange) collected from a mesocosm containing seven fish species. Active filtration eDNA detection results (blue) adopted from Jeunen and von Ammon et al. (*in prep*.). Filled and unfilled circles indicate taxon presence or absence. Circle diameter indicates the number of positive detections within each sampling depth and survey method (maximum 5 replicates for each treatment).

## 3 DISCUSSION

The results presented in this study provide insight into the comparability among four eDNA signal sources, with high similarity between aquatic and sponge eDNA. We provide evidence that both aquatic and sponge eDNA can detect spatially specific eDNA signals, resembling in-field community assemblages, within a 20 m vertical transect across a strong halo-and thermocline. Bivalve gill-tissue dissections, on the other hand, failed to recover eDNA. Also, our results show the feasibility to collect diversity information from ethanol in which filter-feeding specimens are stored, though with more limited success. Furthermore, all eDNA sources were able to detect a greater number of vertebrate taxa than we observed in our diver survey.

### 3.1 Highly similar aquatic and sponge eDNA signals detect community structures induced by water stratification

Environmental DNA signals obtained from water samples and sponge specimens showed high similarity in alpha and beta diversity. Additionally, eDNA signal strength was highly correlated between aquatic and sponge eDNA. Furthermore, both eDNA sources were able to detect the vertical zonation pattern, induced by the presence of a near-permanent halo- and thermocline, previously described through traditional (Grange et al., 1981) and aquatic eDNA metabarcoding surveys (Jeunen et al., 2019b). For example, freshwater (e.g., *Salmo trutta* – brown trout) and terrestrial eDNA signals (e.g., two land birds, *Gallirallus australis* – weka, *Petroica macrocephala* – tomtit) were primarily observed in the LSL, while marine organisms (e.g., *Odax pullus* – greenbone fish, *Meuschenia scaber* – smooth leatherjacket fish, *Odontaspis ferox* – smalltooth sand tiger shark) were mostly detected in the underlying marine layer. Interestingly, sponges might obtain higher spatial resolutions compared to aquatic eDNA in certain instances, as inadvertent water mixing from Niskin sampling resulted in highly abundant eDNA signals from the LSL being observed in aquatic eDNA samples collected at depth, e.g., *Anguilla dieffenbachii* (New Zealand longfin eel), *Galaxias brevipinnis* (climbing galaxias fish), *Gobiomorphus cotidianus* (freshwater common bully), and *Trichosurus vulpecula* (common brushtail possum). Therefore, our results provide conclusive evidence that sponges naturally accumulate eDNA through either their filter-feeding strategy or the entrapment of particulate matter into the tissue matrix via current flow, or both (Mariani et al., 2019; Turon et al., 2020). Hence, the time-intensive pre-processing step of active filtration, currently limiting sample number and volume (Majaneva et al., 2018; Takasaki et al., 2021), could potentially be omitted by incorporating sponge eDNA into the sampling strategy.

The similarity between eDNA sources is particularly striking given the (i) discrepancy in protocol development between eDNA sources (Bowers et al., 2021) and (ii) observed differences between aquatic eDNA and various eDNA sources in previously published research (Koziol et al., 2019; Turner et al., 2015). During the last decade, multiple research groups have optimized laboratory protocols for aquatic eDNA in the marine environment and have shown protocol choice to influence DNA concentration and diversity detection (Bowers et al., 2021; Jeunen et al., 2018; Kawato et al., 2021; Sanches & Schreier, 2020). Using optimized and standardized protocols is, therefore, essential for successful aquatic eDNA monitoring (Bowers et al., 2021). Thus far, protocols have not yet been developed or optimized to extract eDNA from filter-feeding organisms. Given the advances observed in standard filter-based eDNA sampling, it seems probable that additional research on protocol optimization might further increase the efficiency of sponge eDNA surveys, thereby closing the gap of partial overlap in species detection currently observed between aquatic and sponge eDNA.

While showing promise, several limitations and drawbacks might be specifically associated with sponge eDNA surveys, including sample availability and host-related eDNA signals. Within aquatic eDNA surveys, water sampling can be easily achieved under various circumstances, such as at depth (Thomsen et al., 2016) and remote areas (Cowart et al., 2018). Sponge sampling, however, will require accessible dive sites with acceptable diving conditions or destructive sampling (bottom trawling). Additionally, sponges might not be available across all habitats, occur in low abundance, or be considered important keystone species (Bell et al., 2020). Furthermore, sponges are among the less-studied benthic invertebrates with regards to extinction risk and conservation status (Bell et al., 2015). While sponges can regenerate (Henry & Hart, 2005) and non-destructive sponge eDNA sampling has been proposed (Mariani et al., 2019), caution needs to be undertaken to ensure sponge populations are not adversely impacted.

An additional issue might be caused by the presence of host DNA and host-associated communities (Çinar et al., 2019; Turon et al., 2019). The power of aquatic eDNA surveys is the ability to retrieve information about a broad range of taxonomic groups, made possible by the complexity of the eDNA signals contained within water (Ruppert et al., 2019). Thus far, vertebrate diversity has been the focus of sponge eDNA research (Mariani et al., 2019; Turon et al., 2020). The majority of extracted DNA, however, will originate from the host. Therefore, blocking primers will be required when ‘universal’ metabarcoding primers are used, potentially biasing species detection (Wilcox et al., 2014). Furthermore, sponges are known to host species-specific microbiomes (Turon et al., 2019) and invertebrate communities (Çinar et al., 2019) within their tissue matrix. The DNA from these associated communities could potentially hinder the eDNA exploration of these taxonomic groups when eDNA is sourced from sponges.

### 3.2 Bivalve gill-tissue dissections yield no eDNA signals

To increase the flexibility of incorporating filter-feeding organisms into eDNA surveys, eDNA recovery from bivalve gill-tissue dissections was examined. Compared to sponges, bivalves are more abundant and often more readily accessible, occupying a range of habitats where sponges could be scarce or not present (e.g., mudflats: New Zealand cockle – *Austrovenus stutchburyi*), and are sturdier (Riisgård et al., 2013). The abovementioned attributes facilitate manipulation for in-field experimental design (Barthel & Theede, 1986; Honkoop et al., 2003). Additionally, many bivalves are commercially farmed (Tacon, 2020), perhaps creating an opportunity to use such installations as ongoing platforms for biodiversity and biosecurity monitoring. However, as with sponges, co-occurring organisms may complicate analyses. Bivalve whole-genome sequencing efforts have reported high levels of background signal, likely a consequence of parasites and commensal organisms, as well as the accumulation of eDNA through their filter-feeding strategy (Ashby, 2019). Through multiple in-field and mesocosm experiments using two bivalve species (i.e., blue mussel – *Mytilus galloprovincialis*; New Zealand cockle – *Austrovenus stutchburyi*), we provide strong evidence that eDNA is not reliably recovered from bivalve gill-tissue dissections. However, given the huge potential of incorporating bivalves into eDNA surveys, we recommend additional research to be undertaken to investigate if eDNA is accumulated in other tissues, such as the gastro-intestinal tract, targeted in invertebrate diet metabarcoding analyses (Siegenthaler et al., 2019; van der Reis et al., 2018; Yeh et al., 2020).

### 3.3 Diversity detection from preservation mediums

Genetic and genomic information about the specimen has previously been recovered from ethanol and other preservative mediums (Carew et al., 2018; Hahn et al., 2021; Zizka et al., 2018), as cellular material and DNA leaches into the storage buffer. Furthermore, the diversity of bulk invertebrate samples has successfully been sequenced through DNA metabarcoding from the ethanol in which bulk invertebrate samples were stored (Carew et al., 2018; Zizka et al., 2018). Our results show, for the first time, that not only host DNA, but also eDNA can be recovered from ethanol, though with limited success when eDNA is extracted within a month of sampling. The limited diversity recovered from ethanol could potentially be explained by either methodology or time. While our chosen protocol has previously been used to extract DNA from ethanol (Hajibabaei et al., 2012), it is possible that centrifugation, precipitation, or filtration is preferred over evaporation, as these methods could allow for an increased volume of ethanol to be processed. Alternatively, storage time could influence eDNA recovery when (e)DNA leaching is a continuous process until an equilibrium is reached. Hence, further research is required to determine an optimal eDNA extraction protocol and storage conditions for eDNA extraction from ethanol and other preservative mediums (Hahn et al., 2021).

### 3.4 Environmental DNA surveys detect an increased number of taxa compared to diver surveys

All eDNA sources were able to detect a larger number of taxa compared to the diver survey, though three fish observed by divers were not identified by eDNA, i.e., blue cod (*Parapercis colias*; sighted once by diver and noted as ‘rare’ by diver), oblique triplefin (*Forsterygion maryanne*; noted as ‘common’ by diver), and common triplefin (*Forsterygion lapillum*; noted as ‘common’ by diver). These results are in agreement with published research comparing aquatic eDNA to various traditional monitoring methods (Jeunen et al., 2020; Stat et al., 2019; Thomsen et al., 2016). Additionally, diver-observed fish constituted the majority of reads in the eDNA analysis, with a higher proportion of reads assigned to ‘abundant’ fish compared to ‘rare’ fish. These results indicate that abundance information could potentially be obtained from metabarcoding data (Nevers et al., 2018).

False-negative eDNA species detections could potentially be attributed to sensitivity and incomplete reference databases. The two fish species noted as ‘rare’ by diver were either observed in a reduced number of samples (*Pseudolabrus miles* – scarlet wrasse) or not observed (*Parapercis colias* – blue cod), potentially indicating a lack of sensitivity. While limit-of-detection (LOD) experiments identified species-specific eDNA surveys to be highly sensitive (Hunter et al., 2017), LOD is complex to identify for metabarcoding, as multiple parameters will influence amplification efficiency, such as mismatches in primer-binding sites, community diversity, and primer efficiency of co-occurring species (Cristescu & Hebert, 2018; Xiong et al., 2016). Five unique Tripterygiidae (triplefins) eDNA signals were detected, but due to a limited number of reference sequences, these eDNA signals were classified at family, rather than species level. Therefore, both species of triplefins identified by divers could not be identified from eDNA, due to incomplete fish reference databases.

## 4 MATERIALS AND METHODS

### 4.1 Sampling site description

Field sampling was conducted at Bauza Island in Doubtful Sound, a fiord situated in the southwest region of New Zealand (45°17’51.7632”S; 166°55’35.9292”E; Figure 1a) during December 2020. Doubtful Sound is notable for its steep rock wall dropping down to deep water (>400 m), with a sharp, near-permanent halocline, which separates the near-freshwater surface layer (~2-5 m thick) from the underlying full-salinity marine layer (Barker & Russell, 2008).

This persistent low-salinity layer (LSL) on the surface is the result of high rainfall, fiord morphology, and additional freshwater discharge from a large hydroelectric power scheme (Gibbs et al., 2000). This vertical stratification results in strong/distinct vertical zonation of plants and animals in the upper 50 m. Low species diversity is found in the intertidal region and highly diverse assemblages are present below the LSL (Boyle et al., 2001; Grange et al., 1981; Rutger & Wing, 2006).

### 4.2 Contamination prevention

Laboratories and equipment were sterilized by a 10 minute exposure to 10% bleach solution (Prince & Andrus, 1992). Afterwards, all bench space and equipment were wiped with ultrapure water (Ultrapure^™^ DNase/RNase-Free Distilled Water, ThermoFisher Scientific, Cat. NR. 10977015) to remove bleach residue. Sampling bottles were purchased new (2L, HDPE Natural, EPI Plastics) and decontaminated by rinsing twice with ultrapure water, submerging in 10% bleach for 10 minutes, and rinsing twice again with ultrapure water. Negative field controls (500 mL ultrapure water-filled bottle placed amid sampling bottles), filtration controls (vacuum filtration of 500 mL ultrapure water), extraction controls (DNA extraction of 500 µL ultrapure water), and qPCR controls (2 µL ultrapure water) were processed alongside samples to test for contamination during sample handling. Furthermore, all pre-PCR laboratory work was conducted in designated PCR-free eDNA laboratories at Portobello Marine Lab (PML).

### 4.3 Sample collection

#### 4.3.1 Water samples

Five, 2L water samples were collected by Niskin bottles at a depth of 0, 5, 10, and 20 m. Samples were collected following increasing depth to reduce water mixing and contamination. Water samples were transported to the Deep Cove Marine Sciences Field Station on ice in the dark until filtration using a vacuum manifold (Laboport^®^, KNF Neuberger, Inc.) through a 0.45 µm cellulose-nitrate filter (CN; Whatman^™^) within ~1 hour of collection. Filters were rolled up, cut in half, placed in 2 mL DNA LoBind Eppendorf^®^ tubes, and stored in the dark on dry ice during shipment to the University of Otago’s PCR-free eDNA facilities at PML. Samples were stored at −20°C until further processing.

#### 4.3.2 Filter-feeding organisms

After water sample collection, five filter-feeding organisms were collected by divers at each of the aforementioned depths. Due to the lack of sponges in the intertidal zone, blue mussels (*Mytilus galloprovincialis*) were collected at the intertidal region. Predation by echinoderms leads to the disappearance of blue mussels at 5 meters depth (Witman & Grange, 1998), hence, a variety of sponges (phylum: Porifera; Figure 1c) were collected and placed in separate 50 mL falcon tubes. Specimens were transported to the Deep Cove Marine Sciences Field Station on ice in the dark until dissection. Both gills were dissected from blue mussels and a ~5 mm^3^ tissue sample was dissected from sponge specimens. Dissections and specimens were placed in 5 mL DNA LoBind Eppendorf^®^ and 50 mL falcon tubes, respectively. Both 5 mL and 50 mL tubes were filled with 99.8% molecular grade ethanol (Fisher BioReagents^™^, Fisher Scientific) and stored in the dark on dry ice during shipment to the University of Otago’s PCR-free eDNA facilities. Sponges have been taxonomically identified (Figure 1c; Supplement 2) by combining internal and external morphological features (Bergquist, 1998). Samples were stored at −20°C until further processing.

#### 4.3.3 Complementary data collection

Prior to sample collection, water column temperature and salinity were vertically profiled to 20 meter depth by CTD (RBR XR-420 Conductivity, Temperature, Depth Profiler; RBR Ltd, Ottawa, Canada; Figure 1b). Diver visual fish surveys were additionally conducted at each site during filter-feeding specimen collection. Divers also recorded videos via GoPro (GoPro Hero 4 Silver cameras mounted in waterproof housings) along the vertical transect for later verification, and fish species were identified through morphological characteristics to validate fish detections via eDNA signal recovery from water and filter feeder samples. Due to the low visibility in the LSL, diver surveys were only conducted in the marine layer.

#### 4.3.1 Mussel mesocosm experiment

This experiment is, to the best of our knowledge, the first to investigate the use of bivalve gill-tissue for eDNA detection. Additionally, despite the ability of blue mussels to adapt to low-salinity environments with minimal impact on filtration rates (Riisgård et al., 2013), the Doubtful Sound LSL can be seen as a suboptimal environment. Environmental DNA metabarcoding results were, therefore, validated in a mesocosm experiment (Supplement 7). Five blue mussels were collected from the head of the Otago Peninsula, on the east coast of southern New Zealand (45°46’57.8136”S; 170°43’4.7892”E) and transported to University of Otago’s Portobello Marine Laboratory (PML). The blue mussels were placed in a mesocosm containing seven fish species (i.e., spotty wrasse – *Notolabrus celidotus*; banded wrasse – *Notolabrus fucicola*; terakihi – *Nemadactylus macropterus*; trumpeter – *Latris lineata*; olive rockfish – *Acanthoclinus fuscus*; thornfish – *Bovichtus variegatus*; smooth leatherjacket – *Meuschenia scaber*). Mussels were submerged overnight for ~12 hours for acclimatisation purposes. Mussel gill tissues were dissected and placed in 5 mL Eppendorf tubes filled with ethanol and samples were stored at −20°C until further processing. A detailed description of the experimental setup can be found in Supplement 7.

### 4.4 DNA extraction and library preparation

Cellulose-nitrate filters, mussel gill tissue dissections, sponge tissue dissections, and ethanol in which filter-feeding organisms were stored were extracted using Qiagen’s DNeasy Blood & Tissue Kit (Qiagen GmbH, Hilden, Germany) following the manufacturer’s protocol, with sample-specific modifications. A detailed description of DNA extraction protocols can be found in Supplement 8. DNA extracts were stored at −20°C until further processing.

Library preparation followed the protocol described in Berry et al. (2017) and Jeunen et al. (2019). Briefly, samples were amplified using three metabarcoding assays targeting one fragment of the 16S rRNA gene region (fish (16S); Berry et al., 2017) and two fragments of the 12S rRNA gene region (MiFish-U/E; Miya et al., 2015). Prior to library preparation, DNA amplification was optimized using a dilution series (neat, 1/10, 1/100) to identify inhibitors and low-template samples (Murray et al., 2015). Amplification was carried out in duplicate in 25 µL reactions prepared with 1X SensiFAST^™^ SYBR^®^ Lo-ROX Kit (Bioline, Meridian Bioscience), 0.4 µL of the forward and reverse primer (IDT; Integrated DNA Technologies, Australia), and 2 µL of DNA. qPCR conditions for the MiFish-U/E assays included an initial denaturing step at 95°C for 10 min; then 50 cycles of 20 s at 98°C, 15 s at 60°C, 15 s at 72°C; and a final melt curve analysis, while the following conditions were used for the fish (16S) assay: an initial denaturing step at 95°C for 10 min; then 50 cycles of 30 s at 95°C, 30 s at 54°C, 45 s at 72°C; and a final melt curve analysis.

A one-step amplification protocol and fusion primers were used for library building. Fusion primers contained a modified Illumina sequencing adapter, a barcode tag (6-8 bp in length), and the template-specific primer. Each sample was amplified in duplicate following the amplification protocol described above and assigned a unique barcode combination to allow the pooling of samples post-qPCR based on Ct-values, end-point fluorescence and melt curve analysis. Mini pools were size-selected and purified using AMPure XP Beads (BioLabs Inc., USA) prior to quantification on Qubit (Qubit^™^ dsDNA HS Assay Kit, ThermoFisher Scientific) and gel electrophoresis. Pooling of mini pools resulted in a single library. The library was quantified on Qubit and sequencing was performed in-house on an Illumina MiSeq^®^ using a 300-cycle, single-end kit following the manufacturer’s protocols, with 5% of PhiX to minimize issues with low-complexity libraries.

### 4.5 Bioinformatic analysis

Raw sequencing data quality was checked using FastQC v 0.11.7 (Bioinformatics, 2011). Reads were demultiplexed and assigned to samples using cutadapt v 2.10 (Martin, 2011). The assigned amplicons were filtered using the *‘--fastq_filter’* function in VSEARCH v 2.13.3 (Rognes et al., 2016) based on a maximum error of 1.0, maximum length of 230 base pairs, minimum length of 150 base pairs, and removal of sequences containing ambiguous bases. The success of quality filtering was checked in FastQC by comparing reports of FASTQ files before and after the bioinformatic pipeline. Reads passing quality filtering were dereplicated and unique sequences occurring less than 100 times were removed using the *‘--derep_fulllength’* function in VSEARCH. Chimeras were removed using the *‘--uchime3_denovo’* function and unique sequences were clustered at 97% using the *‘--cluster_size’* function in VSEARCH. Finally, an OTU table was generated using the *‘--usearch_global’* function in VSEARCH based on a 97% similarity.

Taxonomy was assigned to each OTU using both BLAST and the Sintax algorithm implemented in VSEARCH. Taxonomy assignments from BLAST results (blastn; megablast; nr/nt database; exclude “uncultured/environmental sample sequences”) were processed by an in-house python script to determine the last common ancestor (LCA; Jeunen et al., 2020). For Sintax taxonomy assignments, custom curated reference databases were generated for each of the three assays using CRABS v.1.0.1 (Jeunen et al., *in prep*.; GitHub: https://github.com/gjeunen/reference_database_creator; Supplement 9). Reference databases were generated by downloading sequences from the NCBI nt and MitoFish databases. In silico PCRs were conducted for each assay, allowing for four mismatches in the primer-binding region. The remaining amplicon sequences were dereplicated to contain unique sequences per species. Further filtering was conducted to exclude environmental and sequences not resolved to species (sequences identified in the database with “.sp”). Additionally, sequences were filtered based on length, ambiguous bases, and those containing missing taxonomic information on Kingdom, Phylum, Class, Order, Family, Genus, and Species level. Final reference databases contained 75,170 sequences covering 38,801 taxa in 11 phyla for the 16S-fish assay and 24,731 sequences covering 15,577 species in six phyla for both MiFish assays. Taxonomy was assigned to each OTU at family level when (i) BLAST and Sintax did not agree on genus-level assignments, (ii) phylogenetic analysis of the family displayed paraphyletic genera for the amplicon region (phylo_build module in CRABS v1.0.1), or (iii) insufficient reference sequences are available to confidently assign taxonomy below family. Genus-level assignment was achieved when (i) BLAST and Sintax agreed on genus or species-level assignment, and (ii) phylogenetic analysis of the family displayed genus-level specificity for the amplicon region.

OTUs receiving identical taxonomic assignments were aligned using MUSCLE (Edgar, 2004), supplemented with reference sequences sharing the same family assignment, and combined into unique taxa. OTUs failing to receive a taxonomic assignment and a positive detection in negative controls were discarded. To avoid issues relating to tag jumping, detections less than 10 reads per sample were discarded (Schnell et al., 2015). Due to a lack of voucher specimens or a local reference database from our sampling site, highest taxonomic resolution was set at genus level for all three metabarcoding assays. Based on existing records of species occurrences within a genus in New Zealand (Ayling, 1987), a “possible species ID” was added to the taxonomic assignment when both Sintax and BLAST achieved high-confidence results for the same species (Supplement 3).

### 4.6 Statistical analysis

Rarefaction curves were generated for each assay separately to assess sequencing coverage using the *‘vegan* v 2.5-7*’* package in R v 4.0.5 (R; http://www.R-project.org). The effects of primer choice on biodiversity detection were visualized using Venn diagrams and measured using pairwise Kulczynski Dissimilarity Indices (KDI; ‘*vegdist*’ function in ‘*vegan’*) at order and taxonomic ID levels (0: equal; 1: unique). Species richness was calculated for each sample and a one-way analysis of variance (ANOVA) and post-hoc Tukey-Kramer test were calculated to determine significant differences in alpha diversity between eDNA sources within and between depths. Species accumulation curves were generated in the *‘BiodiversityR* v 2.13-1*’* package to assess differences in total number of species detected between eDNA sources within each depth. Scatter plots of total number of reads between eDNA sources were generated for all detected species to assess the significance of correlation. A permutational multivariate analysis of variance (PERMANOVA) was used to determine whether eDNA signals differed among depths and eDNA sources. Significant differences in dispersion between groups was tested (PERMDISP) to assess the reliability of PERMANOVA. A principal coordinate analysis (PCoA) was performed to visualize patterns of sample dissimilarity using the Jaccard index (presence-absence transformed data). Analyses were performed using the functions *‘vegdist’, ‘betadisper’* and *‘adonis’* from the *‘vegan’* package. All bioinformatic and statistical scripts can be found in Supplement 10.

## 5 ACKNOWLEDGEMENTS

This work was funded through the University of Otago Research Grant (UORG; UORG2020: #3541 to MD Lamare; Sentinels of change: biodiversity and biosecurity monitoring using environmental DNA from natural samplers).

## 6 AUTHOR CONTRIBUTIONS

GJJ, NJG, and ML designed the Doubtful Sound experiment. Aquatic eDNA sampling at Doubtful Sound was conducted by GJJ, SH, and NJG. Filter-feeder sampling and SCUBA diver surveys were conducted by KE and ML. NP and LU assisted with field work. Laboratory work, including sample pre-processing, DNA extraction, and library preparation, was performed by GJJ, NJG, ML, SF, and JC. Taxonomic identification of sponges through morphology and spicule analysis was conducted by FS. GJJ and UvA designed the mesocosm bivalve experiment. Laboratory work associated with the bivalve mesocosm experiment was performed by GJJ and UvA. GJJ and SF designed the cockle field experiment. Cockle field and laboratory work was conducted by GJJ, SF, POR, and AK. Next-generation sequencing was performed by RD. Bioinformatic and statistical analyses were conducted by GJJ and HC. GJJ wrote the manuscript, with substantial help from JC, FS, UvA, NJG, and ML. All authors contributed to the manuscript writing.

## 7 DATA AVAILABILITY STATEMENT

Raw and demultiplexed sequencing data will be made available on Sequence Read Archive (SRA) upon acceptance of the manuscript being published

