## Supplement 1 for "Assessing the utility of marine filter feeders for environmental DNA (eDNA) biodiversity monitoring"

*Supplemental files:*

Supplement 1: Fish species detected during the 1-hour SCUBA diver survey by three experienced marine scientists in Bauza Island, Doubtful Sound, Fiordland, New Zealand. Abundance of fish was classified as rare (“*”), common (“**”), or abundant (“***”).

| Species name | Common name | Abundance |
| --- | --- | --- |
| *Notolabrus celidotus* | Spotty wrasse | *** |
| *Notolabrus fucicola* | Banded wrasse | ** |
| *Helicolenus percoides* | Jock Stewart | ** |
| *Caesioperca lepidoptera* | Butterfly perch | *** |
| *Parapercis colias* | Blue cod | * |
| *Forsterygion maryannae* | Oblique triple fin | ** |
| *Forsterygion lapillum* | Common triple fin | ** |
| *Pseudolabrus miles* | Scarlet wrasse | * |

Supplement 2: Results of spicules analysis of sponge specimens included in the study. Spicules were photographed under a compound light microscope (Leica Microsystems DM LB) combined with a Canon EOS 70D digital camera. Identified sponge taxa are commonly found in New Zealand coastal waters (Kelly et al., 2009). (a) (b) (c) *Leucettusa lancifera* Dendy, 1924; (d) (e) *Callyspongia* Duchassaing & Michelotti, 1864; (f) (g) *Polymastia hirsuta* Bergquist, 1968; (h) (i) (l) *Strongylacidon conulosum* Bergquist & Fromont, 1988.

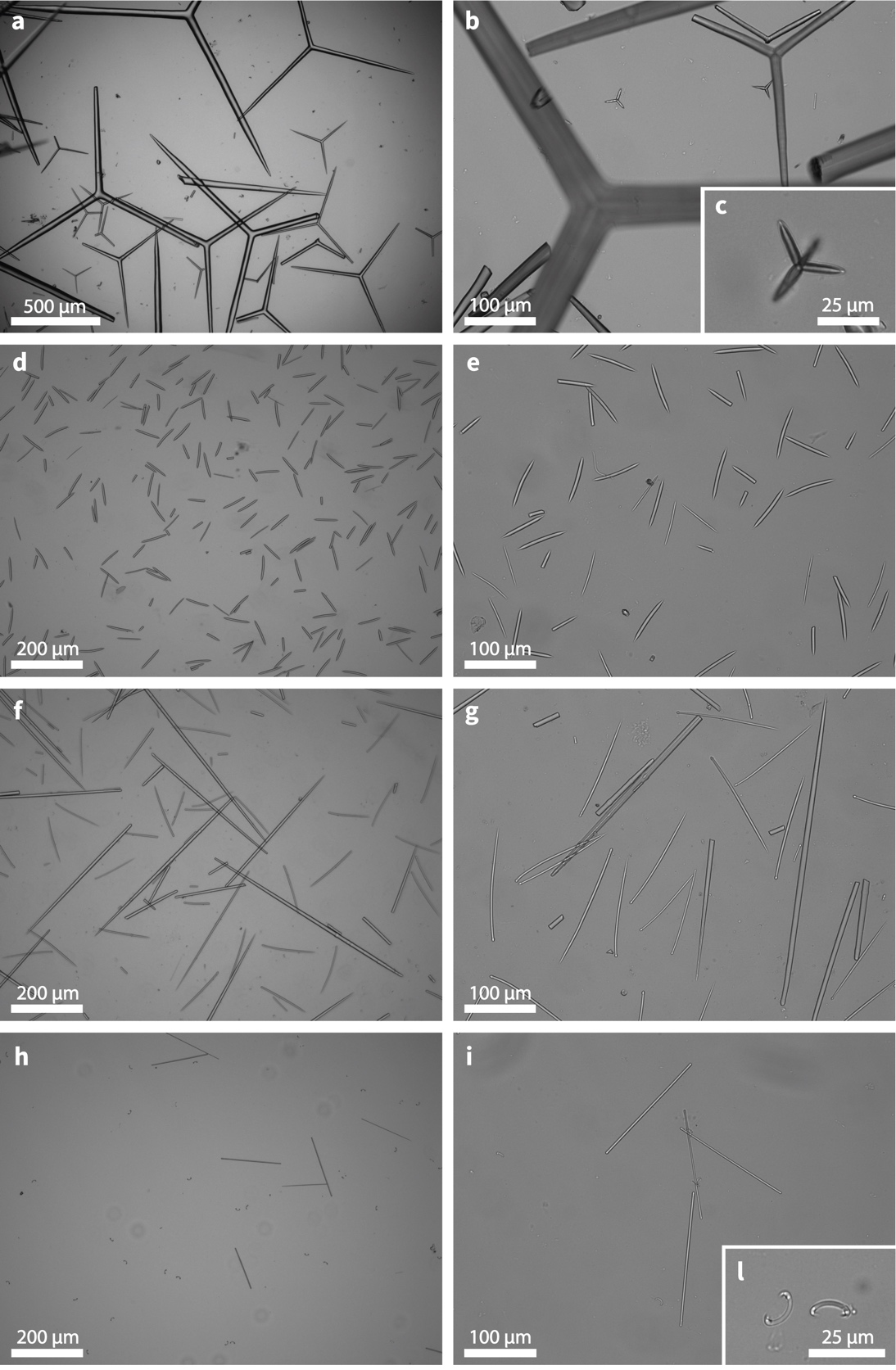

Supplement 3: Environmental DNA detections from the four eDNA sources (aquatic, mussel, sponge, ethanol) for the three metabarcoding assays, i.e., fish (16S), MiFish-E, MiFish-U. A taxonomic lineage is provided for each taxonomic assignment, including domain, phylum, class, order, family, genus, and species. The “taxid” column represents the taxonomic assignment used during statistical analysis and the “potential_species_id” column represents the most likely species assignment for the taxid. Values indicate the number of reads assigned to each taxonomic unit for a given sample. Sample notation follows the following pattern:

BI: Bauza Island

F: aquatic eDNA (filter)

M: mussel eDNA

SP: sponge eDNA

E: ethanol eDNA

First number in between “_”: depth of sample collection

Second number in between “_”: replicate number at a particular site

Data can be found in the “Supplement_3_eDNA_frequency_table.xlsx” file.

Supplement 4: Rarefaction curves for each metabarcoding assay (fish (16S); MiFish-E; MiFish-U) per eDNA source for each sample. Number of taxa are indicated on the y-axis and number of reads on the x-axis. Sample notation follows the abbreviations in Supplement 3. A high resolution figure can be found in file “supplement_4a_rarefaction_curves.eps”.

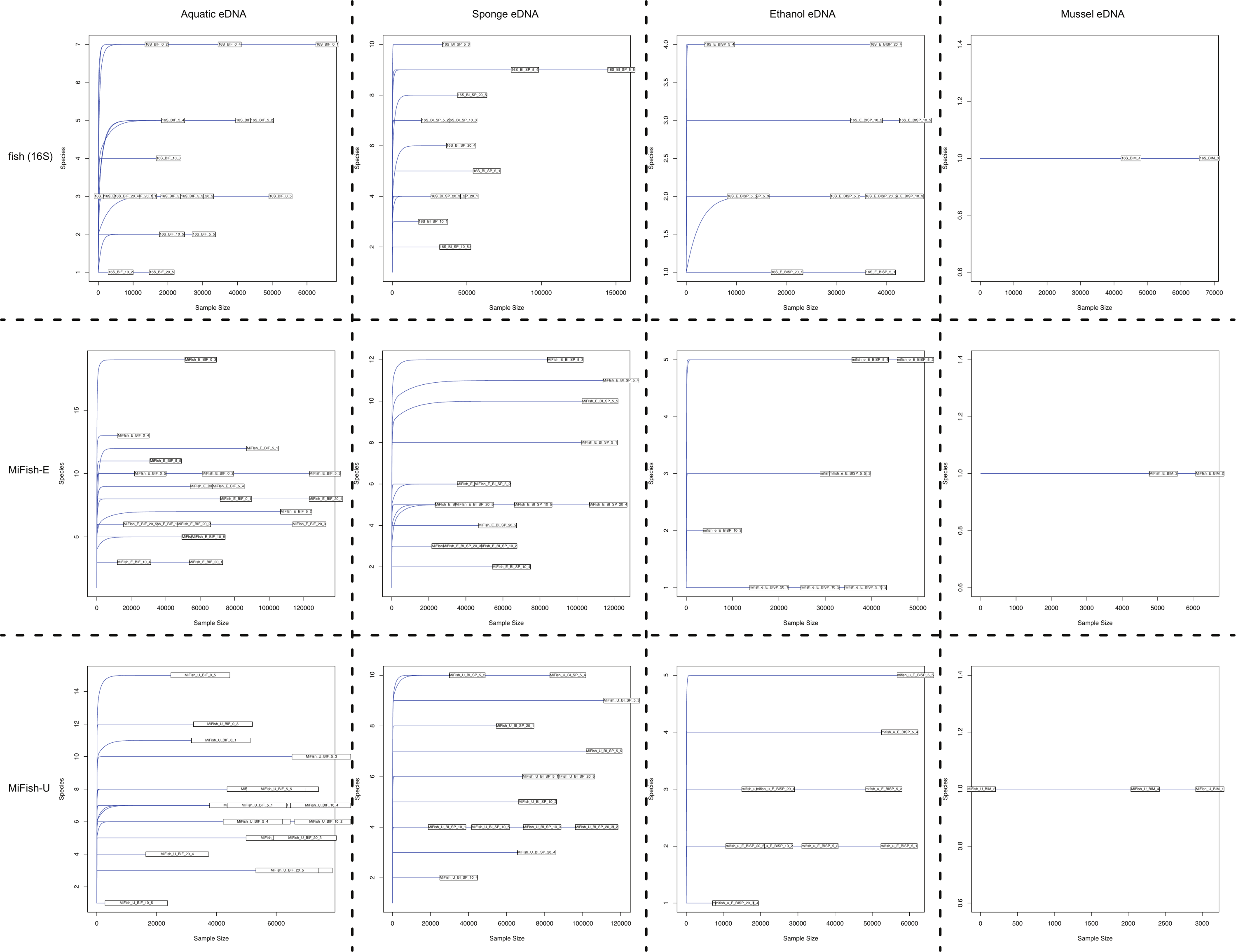

Supplement 5: Taxonomic assignment of Operational Taxonomic Units (OTUs) generated through metabarcoding of 60 samples (aquatic eDNA: 20; mussel eDNA: 5; sponge eDNA: 15; ethanol eDNA (mussel): 5; ethanol eDNA (sponge): 15) using three metabarcoding assays. (a) Number of OTUs assigned to taxonomic ID level. (b) Relative number of total sequences assigned to taxonomic ID level. Shared diversity at (c) order level and (d) taxonomic ID level between the three metabarcoding assays. Percentage of shared diversity is indicated in between brackets. The size of the Venn diagram bubbles is proportional to the total diversity detected by each assay. Overlap without a taxonomic assignment is indicated in grey.

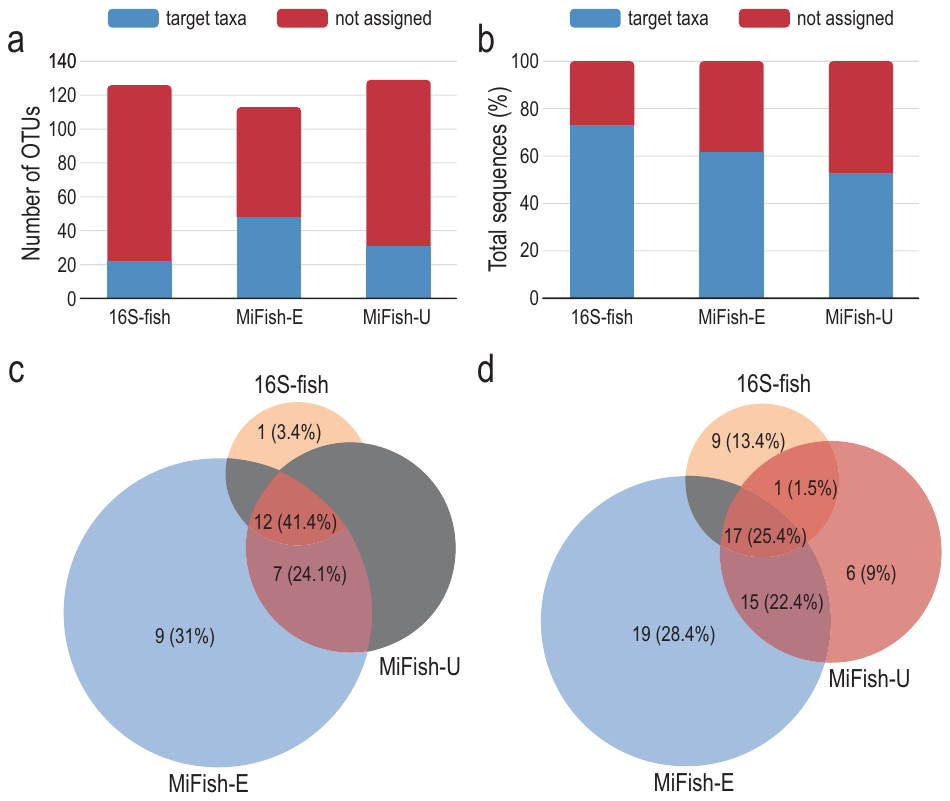

Supplement 6: Experimental setup for the New Zealand cockle (*Austrovenus stutchburyi*) gill tissue eDNA experiment conducted in Aramoana, Dunedin, New Zealand.

**Field and laboratory work**

Cockles (*Austrovenus stutchburyi*) were sampled at low tide on a single day during the summer (January) of 2020, in the mudflats area of Aramoana beach, New Zealand.

Gills were dissected with knife, forceps, and dissecting scissors. Equipment was bleached and rinsed in distilled water between specimens. Gills were placed in a labelled sterile sample vial and kept in a chilly bin with ice packs. Samples were frozen at -20°C until DNA extraction. DNA extraction was performed in a dedicated PCR-free clean room using the DNeasy Blood & Tissue kit (Cat. Nr. 69506) following the manufacturer’s protocols. Samples were amplified following the qPCR protocols described in the main manuscript for all three primer sets. A subset of samples were analysed by gel electrophoresis to confirm amplification and submitted for Sanger sequencing (Genetic Analysis Services, University of Otago) to determine the identity of the amplified products.

**Results and Discussion**

All samples achieved high Ct-values (>40 cycles). Gel electrophoresis only showed positive amplification of the correct amplicon length for samples amplified by MiFish-E (Figure S1). Sanger sequencing of the amplicons revealed bacterial origin of the DNA, indicating unsuccessful amplification of vertebrate 12S rRNA (Table S1). Clean electropherograms were obtained for all sequences (no double peaks), indicating low diversity in the sample. Sequences can be found in files “supplement_6_gill_tissue_rep_1.ab1”, “supplement_6_gill_tissue_rep_2.ab1”, and “supplement_6_blank_rep_1.ab1”.

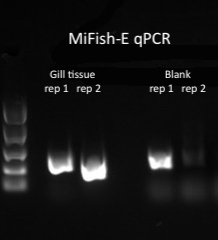

**Figure S1 –** Gel electrophoresis of MiFish-E qPCR assay results; cockle gill tissue samples

show amplified product, such as in these replicates from one particular sample, as well as one of the replicates of the blank extract sample processed during the same extraction protocol.

**Table S1 –** Sanger sequencing BLAST results

| **Sample** | **Query Cover** | **E value** | **% ID** | **BLAST result** | **Species** |
| --- | --- | --- | --- | --- | --- |
| **Gill tissue rep 1** | 95% | 3e-124 | 98.11 | Uncultured bacterium, 16S partial sequence | Uncultured bacterium |
| **Gill Tissue rep 2** | 28% | 1e-28 | 98.73 | Endozoicomonas, 16S partial sequence | *Endozoicomonas* sp*.* |
| **Blank rep 1** | 96% | 7e-121 | 99.19 | Rhizobium sp., 16S partial sequence | *Rhizobium* sp. |

Supplement 7: Experimental setup for the mesocosm experiment (conducted in conjunction with Jeunen & von Ammon et al., *in prep*.).

**Field work and mesocosm setup**

Five blue mussels (*Mytilus galloprovincialis*) were collected on the 21^st^ of April, 2021 from the head of the Otago Peninsula, on the east coast of southern New Zealand (45°46’57.8136”S; 170°43’4.7892”E) and transported to the University of Otago’s Portobello Marine Laboratory (PML). Five blue mussels were placed in a mesocosm containing seven fish species (i.e., spotty wrasse – *Notolabrus celidotus*; banded wrasse – *Notolabrus fucicola*; terakihi – *Nemadactylus macropterus*; trumpeter – *Latris lineata*; olive rockfish – *Acanthoclinus fuscus*; thornfish – *Bovichtus variegatus*; smooth leatherjacket – *Meuschenia scaber*). Mussels were submerged overnight for ~12 hours for acclimatisation purposes.

**Sample collection**

Prior to laboratory work, all bench surfaces and equipment were sterilized in the same manner as discussed in the main manuscript. Mussels were taken out of the mesocosm and opened with a knife. Both gills were dissected using forceps and scalpel. Dissections were placed in 5 mL DNA LoBind Eppendorf^®^ tubes, filled with 99.8% molecular grade ethanol (Fisher BioReagents^TM^, Fisher Scientific). Samples were stored at -20°C until DNA extraction the following day.

Five, 2L surface water samples were collected from the mesocosm in plastic bottles (HDPE Natural, EPI Plastics; acid washed for sterilization purposes) for active filtration. Water samples were filtered over a 0.45 µm cellulose nitrate filter (CN, Whatman^TM^) using a vacuum pump (Laboport^®^, KNF Neuberger, Inc.) and a custom-made manifold. Filters were rolled up, cut in half, each half placed in 2 mL Eppendorf tubes, and stored at -20°C until DNA extraction the following day.

**DNA extraction and further sample processing**

Samples were extracted using the Qiagen DNeasy Blood & Tissue Kit, with slight modifications based on sample type (Supplement 8). Library preparation for DNA extracts followed the methodology described in the main manuscript, with the slight modification that only the fish (16S) primer set was used.

Supplement 8: A detailed, step-by-step description of the DNA extraction protocols for each eDNA source. All samples were processed using the Qiagen DNeasy Blood & Tissue Kit, with slight modifications for each sample type to increase DNA yield.

**Aquatic eDNA:**

- Roll up filters and cut into ca. 1 mm slices.
- Place each half of the filter in a separate 2 mL Eppendorf tube.
- Add 0.5 g of 0.5 mm Zirconia/Silica Beads.
- Add 720 µL ATL buffer and 80 µL Proteinase K.
- Vortex at maximum speed for 10 minutes.
- Incubate samples at 56°C overnight.
- Vortex at maximum speed for 1 minute.
- Centrifuge samples at 6,000 x g for 1 minute.
- Transfer supernatant (600 µL) into a new 2 mL Eppendorf tube.
- Add 600 µL AL buffer.
- Add 600 µL 100% ethanol.
- Vortex at maximum speed for 1 minute.
- Pipet the mixture (620 µL) into a DNeasy Mini spin column placed in a 2 mL collection tube.
- Centrifuge at 6,000 x g for 1 minute.
- Discard the flow-through and collection tube.
- Place the spin column in a new 2 mL collection tube.
- Repeat previous steps until all mixture has passed the spin column.
- Add 500 µL Buffer AW1.
- Centrifuge samples for 1 minute at 6,000 x g.
- Discard the flow-through and collection tube.
- Place the spin column in a new 2 mL collection tube.
- Add 500 µL Buffer AW2.
- Centrifuge samples for 3 minutes at 20,000 x g.
- Discard the flow-through and collection tube.
- Place the spin column in a new 2 mL collection tube.
- Centrifuge samples for 1 minute at 20,000 x g.
- Discard the flow-through and collection tube.
- Transfer the spin column in a 1.5 mL Eppendorf tube with caps removed.
- Add 200 µL AE buffer to the center of the membrane (preheat AE buffer to 56°C).
- Incubate for 1 minute at room temperature.
- Centrifuge samples for 1 minute at 6,000 x g.
- Transfer eluate to a new 1.5 mL Eppendorf tube.
- Store DNA at -20°C.

**Sponge and mussel tissue samples:**

- Store specimens in 99.8% molecular grade ethanol.
- For mussels: place a single gill in a 2 mL Eppendorf tube.
- For sponges: cut a piece of tissue (< 0.3 gram) from the specimen and place in a 2 mL Eppendorf tube.
- Add 720 µL Buffer ATL and 80 µL Proteinase K.
- Vortex at maximum speed for 1 minute.
- Incubate samples at 56°C overnight.
- Vortex at maximum speed for 1 minute.
- Pipette 600 µL into new Eppendorf tube.
- Add 600 µL Buffer AL.
- Vortex at maximum speed for 15 seconds.
- Add 600 µL 100% Ethanol.
- Vortex at maximum speed for 15 seconds.
- Pipette the mixture (620 µL) into a DNeasy Mini spin column placed in a 2 mL collection tube.
- Centrifuge at 6,000 x g for 1 minute.
- Discard the flow-through and collection tube.
- Place the spin column in a new 2 mL collection tube.
- Repeat previous steps until all mixture has passed the spin column.
- Add 500 µL Buffer AW1.
- Centrifuge samples for 1 minute at 6,000 x g.
- Discard the flow-through and collection tube.
- Place the spin column in a new 2 mL collection tube.
- Add 500 µL Buffer AW2.
- Centrifuge samples for 3 minutes at 20,000 x g.
- Discard the flow-through and collection tube.
- Place the spin column in a new 2 mL collection tube.
- Centrifuge samples for 1 minute at 20,000 x g.
- Discard the flow-through and collection tube.
- Transfer the spin column in a 1.5 mL Eppendorf tube with caps removed.
- Add 200 µL AE buffer to the center of the membrane (preheat AE buffer to 56°C).
- Incubate for 1 minute at room temperature.
- Centrifuge samples for 1 minute at 6,000 x g.
- Transfer eluate to a new 1.5 mL Eppendorf tube.
- Store DNA at -20°C.

**Ethanol samples:**

- Pipette 1 mL of ethanol in which a specimen is stored into a 2 mL Eppendorf tube.
- Place with lid open in an oven set at 56°C until fully evaporated.
- Add 180 µL Buffer ATL and 20 µL Proteinase K.
- Vortex at maximum speed for 1 minute.
- Incubate samples at 56°C overnight.
- Vortex at maximum speed for 15 seconds.
- Add 200 µL Buffer AL.
- Vortex at maximum speed for 15 seconds.
- Add 200 µL 100% Ethanol.
- Vortex at maximum speed for 15 seconds.
- Pipette the mixture (600 µL) into a DNeasy Mini spin column placed in a 2 mL collection tube.
- Centrifuge at 6,000 x g for 1 minute.
- Discard the flow-through and collection tube.
- Place the spin column in a new 2 mL collection tube.
- Add 500 µL Buffer AW1.
- Centrifuge samples for 1 minute at 6,000 x g.
- Discard the flow-through and collection tube.
- Place the spin column in a new 2 mL collection tube.
- Add 500 µL Buffer AW2.
- Centrifuge samples for 3 minutes at 20,000 x g.
- Discard the flow-through and collection tube.
- Place the spin column in a new 2 mL collection tube.
- Centrifuge samples for 1 minute at 20,000 x g.
- Discard the flow-through and collection tube.
- Transfer the spin column in a 1.5 mL Eppendorf tube with caps removed.
- Add 200 µL AE buffer to the center of the membrane (preheat AE buffer to 56°C).
- Incubate for 1 minute at room temperature.
- Centrifuge samples for 1 minute at 6,000 x g.
- Transfer eluate to a new 1.5 mL Eppendorf tube.
- Store DNA at -20°C.

Supplement 9: Reference databases generated by CRABS and used by VSEARCH (Sintax) for taxonomy assignment of OTUs. The fish (16S) reference database can be found in file “Supplement_9a_fish_16S_db.tsv” and the MiFish-E/U reference database can be found in file “Supplement_9b_mifish_db.tsv”.

Supplement 10: Bioinformatic and statistical scripts used to process eDNA data. The bioinformatic script can be found in file “supplement_10a_bioinformatic_script.txt” and the statistical analysis R script can be found in file “supplement_10b_statistical_R_script.txt”. Additionally, an updated bioinformatic script in the form of a tutorial (html document with extra information) can be found at <https://github.com/gjeunen/metabarcoding_analysis_tutorial>. This tutorial might deviate from the supplemental file when bioinformatic processing has altered for more recent metabarcoding projects or errors have been discovered in the tutorial.
