## Supplementary figures and images for "Assessing the utility of marine filter feeders for environmental DNA (eDNA) biodiversity monitoring"

### supplement_4a_rarefaction_curves.png

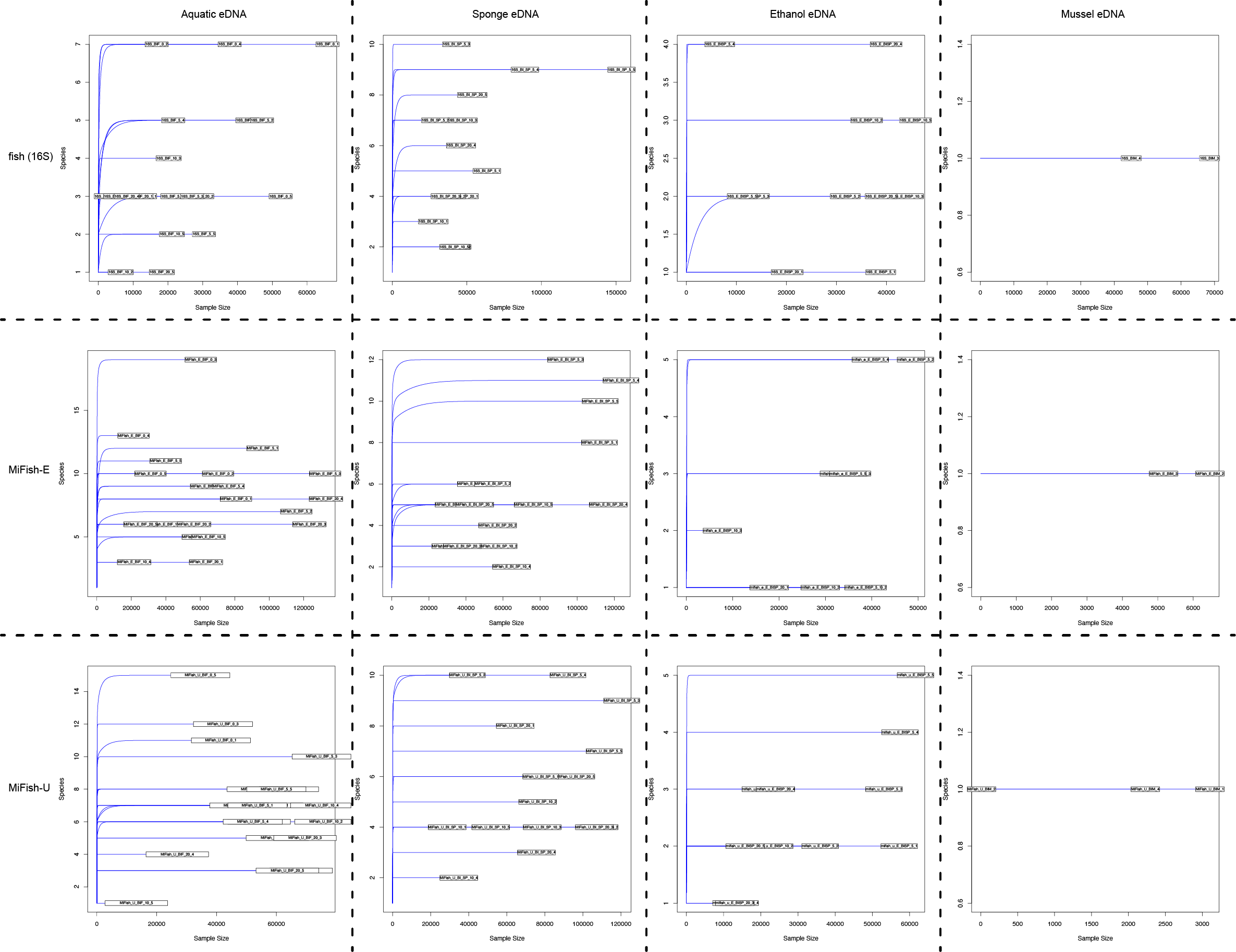
